# Dopamine D2 receptors in nucleus accumbens cholinergic interneurons increase impulsive choice

**DOI:** 10.1101/2023.01.20.524596

**Authors:** Julianna Cavallaro, Jenna Yeisley, Başak Akdoǧan, Joseph Floeder, Peter D. Balsam, Eduardo F. Gallo

## Abstract

Impulsive choice, often characterized by excessive preference for small, short-term rewards over larger, long-term rewards, is a prominent feature of substance use and other neuropsychiatric disorders. The neural mechanisms underlying impulsive choice are not well understood, but growing evidence implicates nucleus accumbens (NAc) dopamine and its actions on dopamine D2 receptors (D2Rs). Because several NAc cell types and afferents express D2Rs, it has been difficult to determine the specific neural mechanisms linking NAc D2Rs to impulsive choice. Of these cell types, cholinergic interneurons (CINs) of the NAc, which express D2Rs, have emerged as key regulators of striatal output and local dopamine release. Despite these relevant functions, whether D2Rs expressed specifically in these neurons contribute to impulsive choice behavior is unknown. Here, we show that D2R upregulation in CINs of the mouse NAc increases impulsive choice as measured in a delay discounting task without affecting reward magnitude sensitivity or interval timing. Conversely, mice lacking D2Rs in CINs showed decreased delay discounting. Furthermore, CIN D2R manipulations did not affect probabilistic discounting, which measures a different form of impulsive choice. Together, these findings suggest that CIN D2Rs regulate impulsive decision-making involving delay costs, providing new insight into the mechanisms by which NAc dopamine influences impulsive behavior.

## INTRODUCTION

Choosing between different reward options requires consideration of their respective costs and benefits. For instance, increasing delay costs can diminish the subjective value of a reward, leading to a preference for immediate rewards[1]. The degree of “discounting” of future rewards, typically measured in humans and animals using delay discounting tasks [2], varies widely among healthy individuals. However, delay discounting can become maladaptive, leading to an excessive bias towards proximal, often less valuable rewards. Indeed, excessive impulsive choice is strongly implicated in substance use disorders (SUDs), attention-deficit hyperactivity disorder (ADHD), schizophrenia and other neuropsychiatric illnesses, as well as in obesity [3-6]. Excessive delay discounting also correlates with risky sexual behavior and overall lack of health monitoring and poor treatment compliance [3,7]. It is not surprising that delay discounting has been proposed as a trans-disease and trans-diagnostic process, reflecting its potential as a candidate treatment target [3,8]. However, the underlying neural substrates and cellular mechanisms remain to be fully understood.

Neuroimaging studies in humans and neuropharmacological and lesion studies in rodents suggest a critical involvement of the nucleus accumbens (NAc) in impulsive choice. Activation of the human ventral striatum, which comprises the NAc, correlates with the subjective value of delayed rewards[9,10]. Lesions of the NAc core subregion in rats reduce preference for large, delayed rewards without affecting sensitivity to reward magnitude [11,12], although partial inactivation of NAc core can decrease delay discounting[13].

The activity of midbrain dopamine neurons has similarly been implicated in delay-based decision-making[14,15]. Given the dense dopaminergic innervation of NAc and the high prevalence of excessive choice impulsivity in disorders that feature ventral striatal dysfunction [3,16], NAc dopamine has been suspected as a key modulator of the region’s role in impulsive choice. While dopamine denervation in NAc failed to alter delay discounting[17], more recent work has demonstrated that phasic dopamine release in NAc *in vivo* encodes delay-related costs and the changing subjective value of rewards [18,19]. Furthermore, optogenetically-evoked NAc dopamine release specifically alters delay-based, but not magnitude-based choices [19].

Various cross-species studies suggest that D2 dopamine receptors (D2Rs) are a critical mediator of dopamine’s actions in these behaviors. Systemic blockade of D2Rs, but not D1Rs, reduces the value of delayed rewards in rats [20], suggesting a causal link between impulsive choice and D2R function. Because such approaches are likely to engage with D2Rs expressed brain-wide, the specific contribution of D2Rs within NAc remains unclear. Positron emission tomography (PET) imaging findings indicate that low D2R availability in the NAc core in rats is correlated with increased impulsivity in a delay discounting task [21,22]. A similar correlation has been reported in ventral striatum of pathological gamblers[16], but whether D2R expression in the NAc leads to impulsive choice involving delayed rewards is unresolved. While neuroimaging and pharmacological studies provide strong support for a role of NAc D2Rs in impulsive choice, they lack the resolution necessary to identify the specific cellular substrates and mechanisms of dopamine-D2R actions. This is especially relevant in the NAc, where D2Rs are widely expressed in spiny projection neurons (SPNs), cholinergic interneurons (CINs), and in presynaptic dopaminergic and glutamatergic axon terminals and can have distinct cellular signaling outcomes[23]. Moreover, given the greater relative abundance of SPNs, observations made with global approaches may obscure important cellular and behavioral contributions of D2Rs expressed in sparser neuronal populations.

Among these, CINs — the main source of acetylcholine in striatum [24,25] — emerge as an intriguing candidate substrate for impulsive behavior. First, CINs influence striatal output by modulating cortico-striatal plasticity in SPNs [26,27], which are thought to play key roles in action selection and reward valuation [28,29]. Second, CINs not only powerfully control local dopamine release [30-32], but their cue-evoked firing activity and acetylcholine release is, in turn, sensitive to dopamine actions on D2Rs [33-35]. Third, recent work involving systemic administration of cholinergic receptor agonists and antagonists has suggested a complex involvement of acetylcholine in delay and probabilistic discounting tasks [36-39]. However, whether D2R function in NAc CINs is critical to impulsive choice has not been investigated.

We recently reported that selective D2R overexpression in NAc CINs impairs learning to suppress responding in a Go/No-Go task when an inhibitory response was required [34]. While this finding is consistent with increased action impulsivity, it is unknown whether NAc CIN D2Rs contribute to impulsive choice behaviors. To this end, we used region and cell type-selective approaches to alter D2R expression in NAc CINs. We found that higher D2R levels in these neurons increase impulsive choice, but only when it involved temporal, but not probabilistic costs. This effect was not associated with altered sensitivity to reward magnitude or impairments in timing. These findings suggest a novel interaction between NAc dopamine and acetylcholine in mediating delay-based impulsive choice.

## MATERIALS AND METHODS

### Mice

Adult male and female hemizygous ChAT-Cre mice (B6.FVB(Cg)-Tg(Chat-cre)GM60Gsat/Mmucd, GENSAT; MMRRC stock no. 030869-UCD)[40], backcrossed > ten generations to C57BL/6J background, were used in D2R overexpression experiments. For knockout experiments, mice were generated from crosses of hemizygous ChAT-IRES-Cre (*ChAT*^*tm1(cre)Lowl*^*/MwarJ*; JAX stock no. 031661) to Drd2^*loxP/loxP*^ (*Drd2*^*tm1*.*1Mrub*^*/J*, JAX stock no. 020631)[41] mice. The ChAT-IRES-Cre/Drd2^*loxP*^ progeny were then crossed to Drd2^*loxP/loxP*^ to generate ChAT-IRES-Cre/Drd2^*loxP/loxP*^ (CIN-D2KO) and Drd2^*loxP/loxP*^(WT controls). Mice were housed in groups of 3-5 per cage on a 12-h light/dark cycle, and all experiments were conducted during the light cycle. All experimental procedures were performed in accordance with NIH guidelines and were approved by Institutional Animal Care and Use Committees of Fordham University and of the New York State Psychiatric Institute.

### Surgical Procedures

Mice (> 10 weeks) underwent stereotaxic surgical procedures under ketamine-induced anesthesia in which they received Cre-dependent double-inverted open-reading frame (DIO) adeno-associated virus (AAVs) bilaterally into the NAc (400 nL/side). Infusions were done using Bregma-based coordinates: AP, + 1.70 mm; ML, ± 1.20 mm; DV, −4.1 mm (from dura) at a rate of 20 nl/s (20 pulses, 5 min). Viruses used: AAV2/9-EF1a-DIO-D2R(S)-P2A-EGFP[34]; AAV2/5-hSyn-DIO-EGFP (Addgene # 50457-AAV5) or AAV2/9-Syn-DIO-EGFP (Addgene # 100043-AAV9). Assignment of AAV was counterbalanced for sex, age, and home cage origin. Behavior experiments began at least 4 weeks following viral infusions.

### Operant Apparatus

Mice were run in sixteen operant chambers (model ENV-307w, Med-Associates, St. Albans, VT) each inside of a light- and sound-attenuating cabinet. The chamber interior (22 × 18 × 13 cm) was equipped with a liquid dipper resting in a feeder trough that administered a drop of evaporated milk when raised. The feeder trough was centered on one wall in the chamber, and head entries into this trough were measured via an infrared photocell detector. Retractable levers were located on either side of the feeder trough, and each had an LED light above it. The flooring of the chamber consisted of metal rods placed 0.87-cm apart. An audio speaker (ENV-324W) was used to deliver tone stimulus (90 dB, 2500 Hz). The chamber was illuminated by a house light mounted on the wall opposite the trough during all sessions. The experimental protocols were controlled via Med-PC computer interface and Med-PC IV or V software. Behavioral events were recorded with a temporal resolution of 10 ms.

### Dipper and lever press training

For behavior experiments, mice were food-restricted and maintained at 85-90% of their baseline body weight; water was available *ad libitum*. For dipper training, 20 dipper presentations were separated by a variable inter-trial interval (ITI) and ended after 20 rewards were retrieved or after 30 min had elapsed. Mice moved to the second phase of dipper training if head entries were made in 20 dipper presentations in one session. In this training session, criterion was achieved when mice retrieved 30 of 30 dippers[42]. Lever press training was done using a fixed ratio-1 (FR-1) schedule, where each press led to 1 reward. Dippers were raised for 5 s. FR-1 training was done in 2 sessions per day (one for each lever; order alternated each day). The session ended following 20 reinforcers earned, or after 30 min. Sessions were repeated daily until all mice earned 20 reinforcers on each lever.

### Delay Discounting and Probability Discounting Tasks

After successful completion of the criteria for trough and lever press training, mice were trained on a delay discounting procedure adapted for mice[43]. Initial lever preference was first determined from three sessions of two-lever FR-1. The less preferred lever was then assigned the large reward (3 milk dippers given in succession) and the preferred lever was assigned the small reward (1 dipper). Delay discounting sessions began with 10 forced choice trials. In forced choice trials, one of the lever lights appeared for 5 s before the extension of its associated lever; only one lever was presented in forced trials. In forced “delayed” trials, pressing of the corresponding lever led to the large reward after a delay. In “forced immediate” trials, responding on the alternate lever led to a small reward with no delay. The order of forced trials was randomized. Both levers were rewarded on a FR-1 schedule and retracted following a press. In the remaining 20 “free choice” trials, both levers were extended following 5-s presentation of both lever lights. New trials began following a variable ITI (mean = 29 s). For the first 14 daily sessions, the delay to the small and large rewards was set to 0 s, to assess preference for reward size. These sessions were followed by sessions in which the delay to large reward was increased across sessions, while the delay to the small reward remained at 0 s. Time delays (2, 4, 6, 8, and 10 s) to the large reward following a lever press were presented in separate sessions (3 sessions for each delay) in ascending order[43].

Following successful completion of trough and lever press training, separate cohorts were trained on the probabilistic discounting task, where the probability to receive the large reward was progressively decreased across sessions[43]. The initial training phase for this task was identical to that of the delay discounting task, where pressing either lever resulted in a small or large guaranteed reward with no delay. After 14 days of this phase, the probability to receive the large, but not the small, reward was decreased (80, 60, 50, 40, 33, 20%) across sessions. Each probability was presented for 3 consecutive days.

### Temporal Discrimination

A different cohort of mice underwent training on a temporal discrimination task [44]. Following initial dipper training, mice received FR1 lever training, in which a reward was given immediately following a lever press that occurred within the first 30 s of lever presentation or if no press was made for 30 s. In both cases, the lever retracted upon dipper presentation. Trials occurred on a variable ITI (mean = 45 s). These FR1/FT30 sessions lasted until mice earned 30 rewards in 2 consecutive sessions. For discrimination training, mice learned to press one of the two levers after a 2-s tone (“short”) and the other following an 8-s tone (“long”) to earn a reward. The first two days of training involved presenting only one sample duration and its corresponding lever, followed by two days where half the trials were short-tone and half long-tone with single lever presentations. For the next 3 days, 50% of the trials presented either sample durations followed by extension of both levers (choice response trials), while the other 50% of trials presented a single lever associated with the correct categorization of the sample duration (forced response trials). Mice were then trained on a 75% choice response session for 3 days, followed by 15 100% choice response sessions. The durations of the short and long tones were increased from 2 to 6 s and from 8 to 24 s and original lever assignments were maintained. Sixteen additional sessions of this type were conducted.

### Peak Interval Training

Following the temporal discrimination task, the same mice received one additional FR1 session before commencing fixed interval training. Here, lever presses were only reinforced until after a fixed interval (timed relative to lever extension) had elapsed. Each reinforcement was followed by a variable inter-trial interval (mean = 30 s) during which the lever remained retracted. New trials were signaled by lever extension. Mice began with FI-4 s session and proceeded to longer interval sessions after earning ≥ 40 rewards in each session. The FI durations were 4, 8, 16, and 24 s. In peak interval training, a target interval of 24 s was used as described [45]. Each training session consisted of FI-24 and peak trials. In peak trials, the lever was extended for 72-96 s, but lever presses had no consequences. Initially, mice were presented with a random combination of 48 FI-24 s trials and 12 peak trials. Once they earned 40 rewards, each session consisted of 36 FI-24 s trials and 24 peak trials. Sessions ended after 90 min or when mice completed 60 trials.

### Open Field

A separate cohort of mice was tested in open field boxes equipped with infrared photobeams to measure locomotor activity (Med Associates, St. Albans, VT). Data were acquired using Kinder Scientific Motor Monitor software (Poway, CA) and expressed as total distance traveled (m) over 90 min.

### Data analysis

In discounting tasks, the percent of free choice trials in which the large reward option was chosen was determined for each delay by dividing the number of presses on the large reward lever by the total number of presses. Data were expressed as the average of the last 2 sessions at each delay or probability. For temporal discrimination, data was expressed as the proportion of correct responses made on a given lever based on the sample duration presented. Peak interval data used for analysis was averaged across the last 5 of 11 sessions. Sample sizes were determined by performing statistical power analyses based on effect sizes observed in preliminary data or on similar work in the literature. Statistical analyses were performed using GraphPad Prism 9 (GraphPad) and MATLAB (MathWorks). Data are generally expressed as mean ± standard error of the mean (SEM). Unpaired two-tailed Student’s t-tests were used to compare two-group data. Multiple comparisons were evaluated by two-way repeated measures ANOVA, when appropriate. A p-value of < 0.05 was considered statistically significant.

Investigators were blinded to the genotype throughout behavioral assays and data analysis.

## RESULTS

### D2R upregulation in NAc CINs increases delay discounting

To determine whether increased D2R levels in NAc CINs contribute to impulsive choice, we first delivered Cre-dependent adeno-associated viruses (AAV) expressing D2R-EGFP or EGFP into the NAc of adult ChAT-Cre mice (8 mice/group). We previously demonstrated that this manipulation leads to stable and selective overexpression of D2Rs in CINs in this region (2.4-fold over EGFP)[34]. Four weeks after surgery, mice were trained on a delay discounting task adapted to mice [43,46] that measures the choice between pressing one lever to obtain a small, immediate reward or pressing another lever to obtain a three times larger reward that is presented after increasing delays **(Figure 1A)**. Each session started with 10 “forced” trials in which only one lever was presented (five trials on each lever, randomly distributed), followed by 20 trials in which both “small” and “large” levers were presented simultaneously (“free choice” trials).

**Figure 1.**
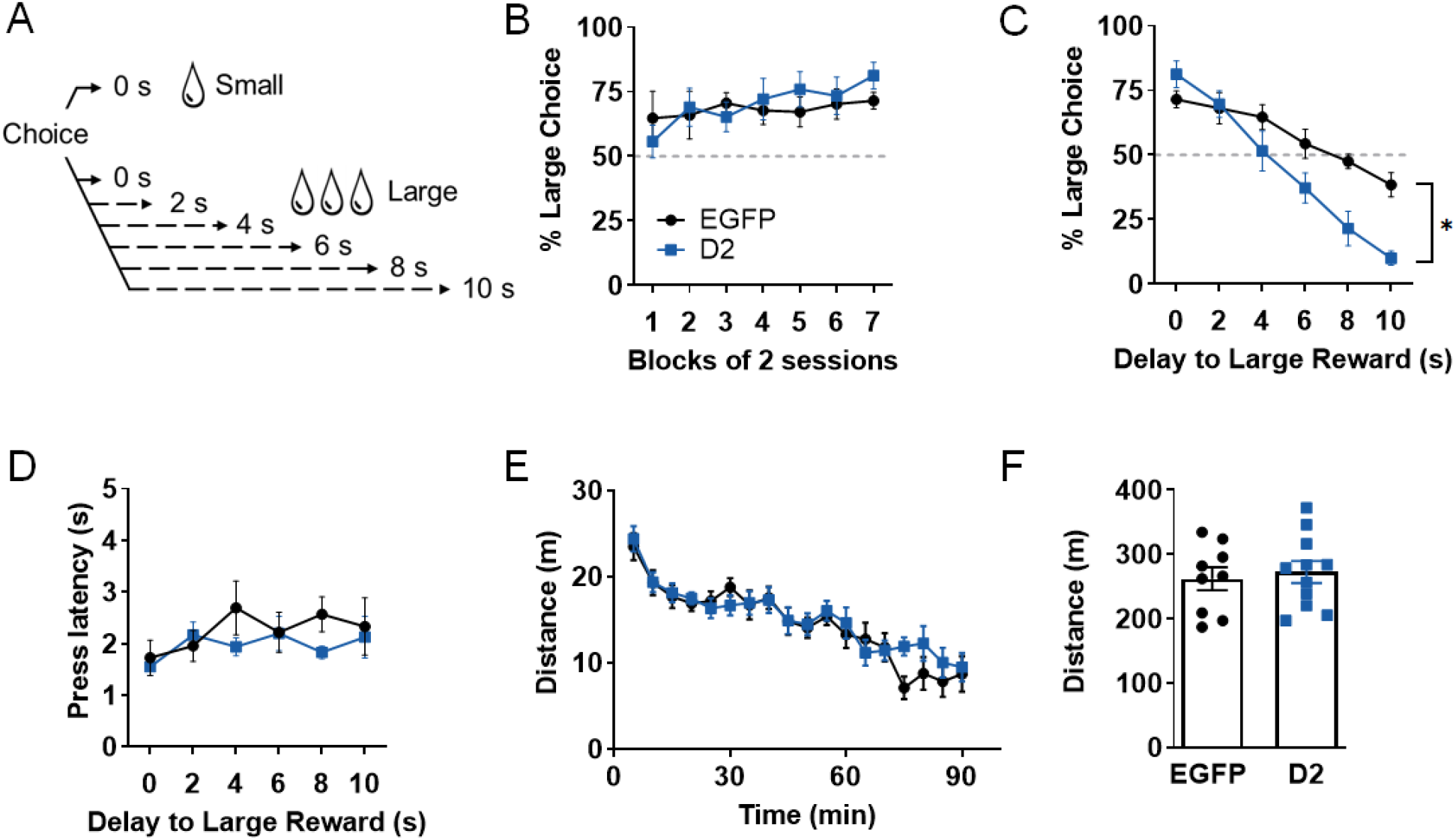
D2R upregulation in NAc CINs increases delay discounting. **A**. Schematic illustration of delay discounting task. On free choice trials, two lever options are presented, each leading to a small or a large reward. The delay to the large reward is progressively increased across sessions (0 – 10 s), while the small reward is given with no delay. **B**. In the absence of delays to either reward, EGFP and D2R-overexpressing mice similarly increased preference for the large reward option after 14 training sessions (shown here as blocks of 2 sessions). **C**. With increasing delays to the large reward, both groups showed discounting, an effect that was significantly greater following CIN D2R upregulation. *p < 0.0001 denotes significant virus x delay interaction, n = 8 mice/group. **D**. In the same mice, latency to make a choice following lever extension in free choice trials was not altered by D2R upregulation. **E, F**. No significant changes were observed in distance traveled in a 90-min open field session (EGFP n = 9; D2 n = 11).

We first assessed the percent choices made on the “large” lever in the absence of delays to either lever over the course of two weeks **(Figure 1B)**. We found that both groups increased their preference for the large reward option with continued training (F _(6, 84)_ = 2.608, p = 0.0229). However, we found no main effect of virus (F _(1, 14)_ = 0.07516, p = 0.7880), suggesting that D2R upregulation does not alter the sensitivity to relative differences in reward size. Following this initial phase, mice experienced increasing delays to the large reward following a lever press, while the small reward continued to be delivered without delay. Delays to the large reward (0, 2, 4, 6, 8, 10 s) were presented in separate sessions for 3 days each. As shown in **Figure 1C**, both groups showed discounting of the large reward as delays increased, as shown by their decreasing choice of the large reward (delay effect: F _(5, 70)_ = 43.47, p < 0.0001). This discounting, however, was significantly steeper in D2R-overexpressing mice compared to controls (virus x delay interaction: F _(5, 70)_ = 6.13, p < 0.0001), suggesting that D2R upregulation in NAc CINs increases impulsive choice. Furthermore, neither the latency to make a selection in choice trials (**Figure 1D**) nor the distance traveled in an open field were altered by this manipulation (**Fig 1E,F**), suggesting that the increased impulsive choice is unlikely to be due to general alterations in motivation or locomotor activity.

### D2R upregulation in NAc CINs does not alter probabilistic discounting

Because rewards obtained after long delays in our delay discounting paradigm could be perceived as being less certain than those obtained after short delays [47,48], it is possible that the effect of CIN D2R upregulation on delay discounting is largely driven by an enhanced intolerance to reward uncertainty. We addressed this issue using a probabilistic discounting task in which subjects must choose between a small, certain reward or larger reward delivered with decreasing probability (**Figure 2A**). While performance in probability discounting and delay discounting depends on an intact NAc core, these two forms of impulsive choice behavior are generally considered dissociable processes[11,47].

**Figure 2.**
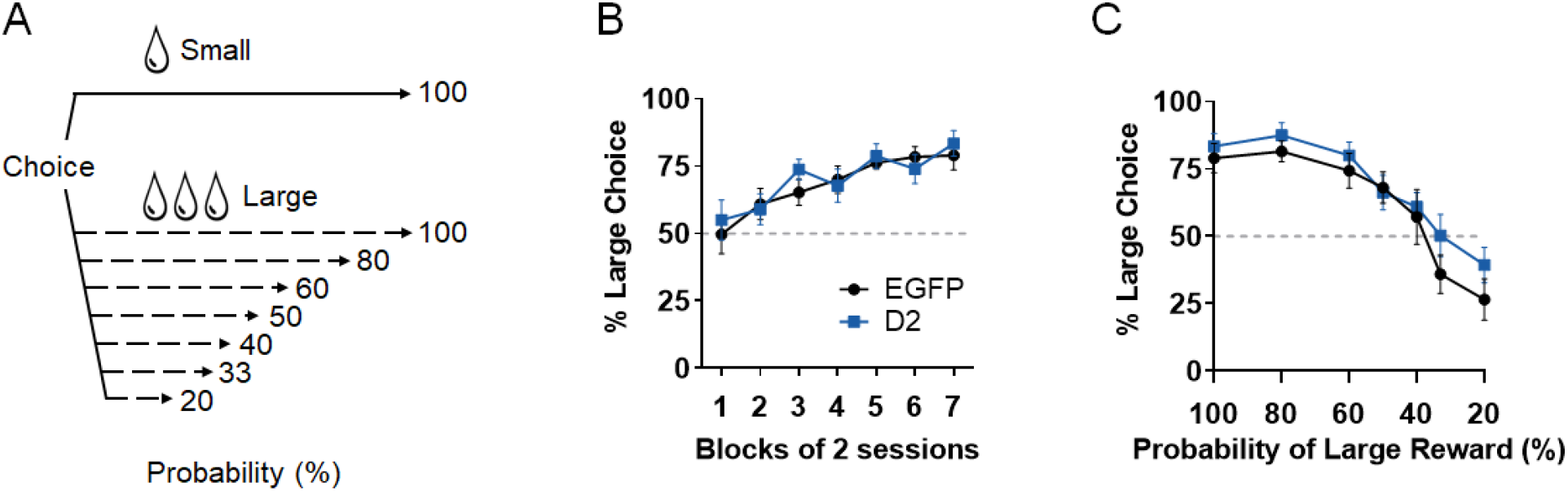
D2R upregulation in NAc CINs does not alter probabilistic discounting. **A**. Schematic illustration of probabilistic discounting task. On free choice trials, two lever options are presented, each leading to a small or a large reward. The probability of receiving the large reward is progressively decreased across sessions (100 – 20%), while the small reward is always given. **B**. With increased training, both EGFP and D2R-overexpressing mice increased preference for the large reward when both options were 100% certain. **C**. Both groups discounted the large reward option as a function of decreased probability of its receipt, but this effect was not significantly different following D2R upregulation (n = 8 mice/group).

We used a variation of the probabilistic discounting paradigm[43] in a different cohort of ChAT-Cre mice overexpressing either D2Rs or EGFP in NAc CINs (n = 8 mice/group). Initial training for this task was identical to the delay discounting task, involving two levers that, when pressed, led to a large or a small reward with 100% probability and 0-s delay. Mice from both groups similarly increased their preference for the large reward across these sessions (F _(6, 84)_ = 17.85, p < 0.0001) (**Figure 2B**). Following this phase, the probability of small reward remained at 100%, while the large reward progressively decreased across sessions (80, 60, 50, 40, 33, 20%), and % preference for the large, uncertain reward was determined. Two-way RM ANOVA indicates that while there was a main effect of probability on discounting (decreased large certain reward choices; F _(6, 84)_ = 38.39, p < 0.0001), there was no significant effect of D2R upregulation (virus x probability: F _(6, 84)_ = 0.7642, p = 0.6001) (**Figure 2C**). These results contrast with our delay discounting findings, suggesting that augmented CIN D2R expression preferentially increases impulsive choice behavior involving delayed reinforcement.

### Genetic inactivation of CIN D2Rs decreases delay discounting but does not affect probabilistic discounting

To determine whether D2Rs in CINs are required for impulsive choice, we selectively deleted the D2R gene in CINs using a ChAT-IRES-Cre x Drd2^*loxP/loxP*^ (CIN-D2KO) mouse line that has been well characterized in several studies of striatal CIN function[35,49-51]. As shown in **Figure 3A**, CIN-D2KO did not differ from Drd2^*loxP/loxP*^ control mice in increasing their preference for the large reward in delay discounting 0 s phase (delay effect: F _(6, 91)_ = 5.112, p < 0.0001; genotype x delay: F _(6, 91)_ = 0.3373, p = 0.9155), suggesting that a lack of CIN D2Rs does not alter reward magnitude sensitivity. However, compared to Drd2^*loxP/loxP*^ mice, CIN-D2KO mice showed decreased delay discounting as evidenced by the greater choice of the large reward option at longer delays (delay effect: F _(5, 65)_ = 72.56, p < 0.0001; genotype x delay: F _(5, 84)_ = 4.756, p = 0.0009) **(Figure 3B)**. In contrast, probability discounting was not affected (**Figure 3C**) (probability effect: F _(6, 72)_ = 14.26, p < 0.0001; genotype x probability: F _(6, 72)_ = 0.9263, p = 0.4814). These findings suggest that CIN D2Rs are required for appropriate discounting of delayed rewards, but do not play a role in discounting of probabilistic rewards.

**Figure 3.**
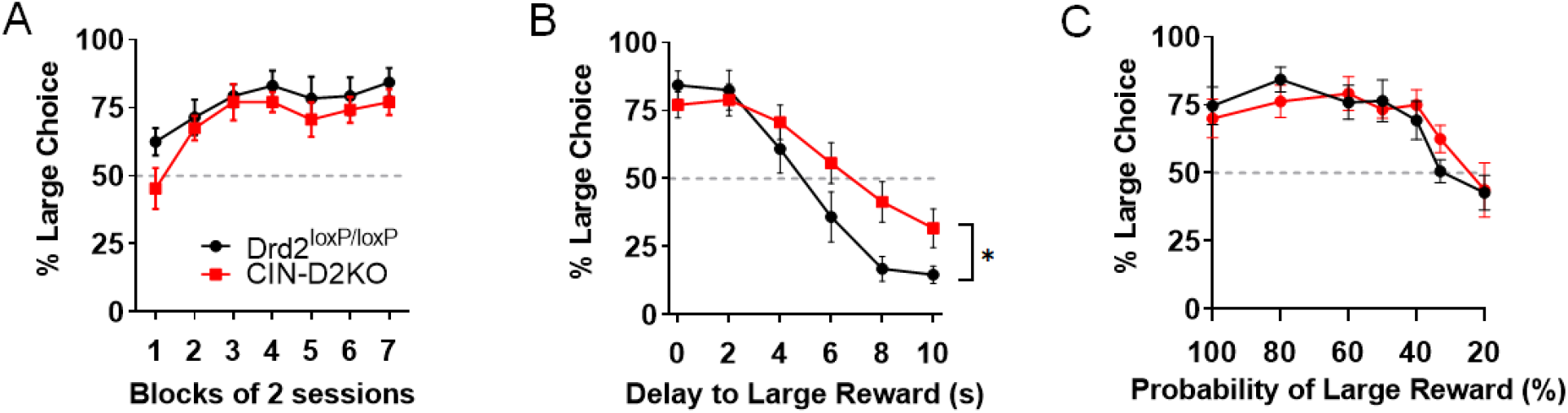
Lack of CIN D2Rs decreases delay, but not probabilistic, discounting. **A**. ChAT-IRES-Cre mice were crossed to Drd2^*loxP/loxP*^ mice to obtain mice lacking D2Rs in CINs (CIN-D2KO). CIN-D2KO and control Drd2^*loxP/loxP*^ mice increased preference for the large reward with training in the absence of delays to either reward option. **B**. Compared to Drd2^*loxP/loxP*^ mice, CIN-D2KO mice showed greater preference for the large reward at longer delays (decreased delay discounting). *p < 0.001 denotes significant genotype x delay interaction (control, n = 8; CIN-D2KO, n = 7). **C**. No significant group differences were observed in probabilistic discounting using a different cohort of mice (control, n = 8; CIN-D2KO, n = 6).

### D2R upregulation in NAc CINs does not alter timing

The ability to accurately represent the time it takes to receive a reward following a press is a key behavioral sub-component in delay discounting tasks[52]. Thus, it is conceivable that CIN D2R upregulation results in an overestimation of time intervals, thereby reducing tolerance of delays compared to controls. To test this hypothesis, we first used a temporal discrimination task to determine whether D2R upregulation altered the ability to correctly categorize two auditory tones of different durations as short or long[44]. Specifically, D2R-overexpressing or control mice were presented either a 2-s (“short”) or an 8-s (“long”) tone, followed by presentation of two choice levers (**Figures 4A**,**B**). A single response on one of the levers was rewarded after the “short” tone, while one press on the other lever was rewarded after the “long” tone. The mean proportion of correct responses during “short” or “long” trials across test sessions was not affected by D2R upregulation (short: F _(1, 13)_ = 0.09168, p = 0.7668; long: F _(1, 13)_ = 0.02325, p = 0.8812; n = 8 EGFP, 7 D2). To determine whether there might be distortions that are specific to particular time ranges [44], we proportionally increased the tone durations for the previously defined “short” and “long” levers to 6 s and 24 s in the same animals (**Figures 4C**). Following the switch to the 6-s (“short”) versus 24-s (“long”) sessions, mice initially exhibited near chance performance in both trial types, likely due to the similarity between 6-s tone and the 8-s tone, which was previously mapped to the “long” lever. A two-way RM ANOVA showed that discriminative performance improved in both groups of mice with training (short: F _(7, 91)_ = 10.67, p < 0.0001; long: F _(7, 91)_ = 10.87, p < 0.0001), without a significant virus x session block interaction (short: F _(7, 91)_ = 0.2038, p = 0.9839; long: F _(7, 91)_ = 0.6794, p = 0.6794).

**Figure 4.**
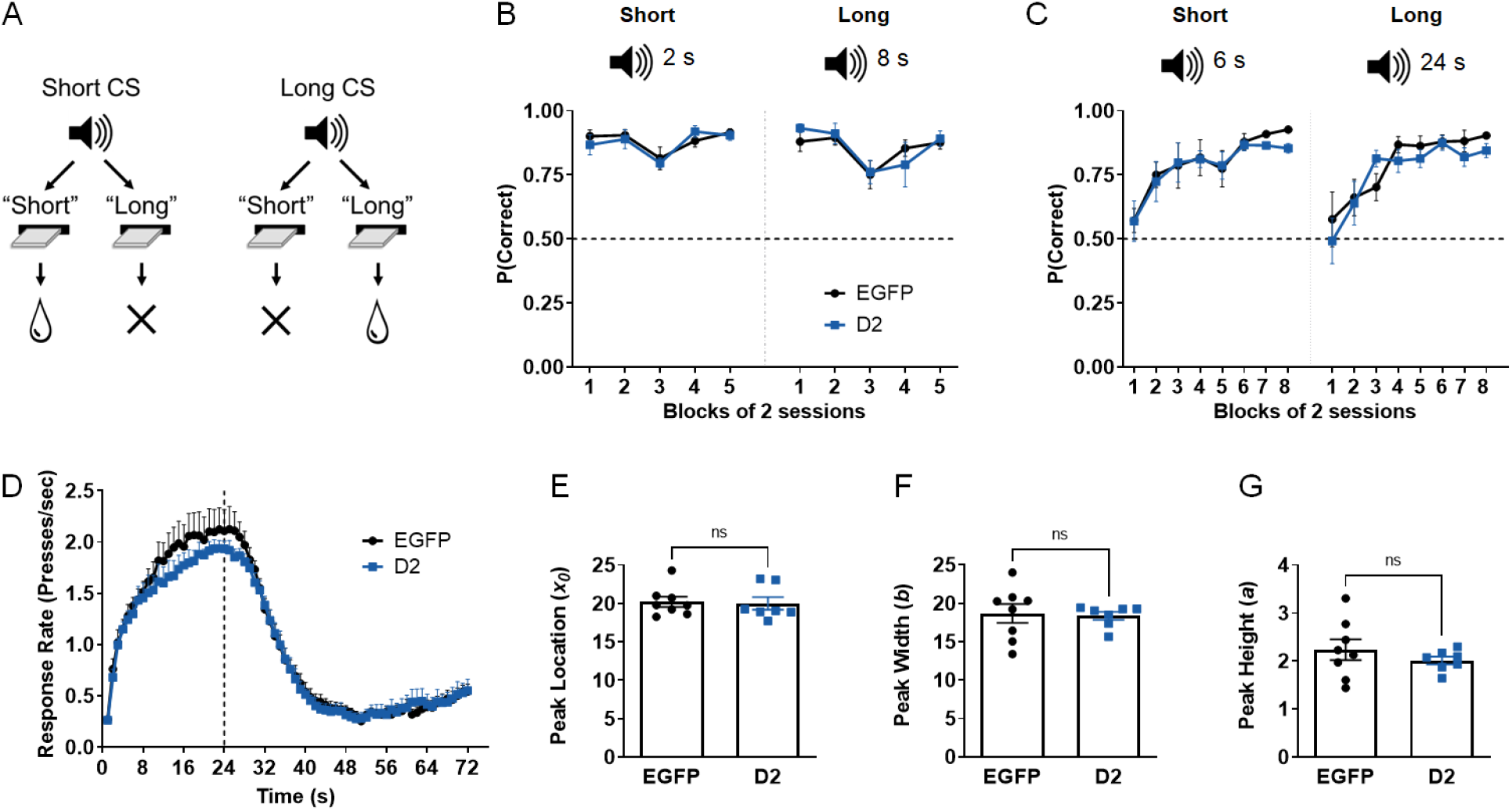
CIN D2R upregulation in NAc does not alter timing. **A**. Schematic representation of the temporal discrimination task. In each session, a single response on one of two lever options is rewarded based on the duration of the sample auditory conditioned stimulus (CS). Two tone durations are presented in each session. **B**. Mean proportion of correct responses on the corresponding lever in 2-s tone trials (short, *left*) or 8-s tone trials (long, *right*) across blocks of 2 sessions. **C**. The duration of tones was proportionally increased to 6 s (short, *left*) and 24 s (long, *right*). CIN D2R upregulation did not alter discrimination of tone durations in either combination. **D** Mean lever press rate during peak trials in the final five sessions of the 24-s peak interval task. **E-G**. Mean best-fitting parameters (derived from fitting to Gaussian function) for peak location (timing accuracy), peak width (timing precision), or peak height (peak response rate). No significant differences were observed in any of these parameters (EGFP, n = 8; D2, n = 7).

Using the same mice, we then examined whether D2R upregulation impacted the precision and accuracy of timing using a peak interval task. In this procedure, mice initially learn that lever responses are only rewarded if they occur after a fixed interval of 24 s (FI-24 s). Peak trials, in which the lever is extended but responses are not rewarded, are then introduced randomly with FI trials [45]. Responding (lever presses/sec) during these peak probe trials was used to examine how accurately and precisely mice time a 24 s interval. **Figure 4D** shows the response rate during peak trials as a function of time in session, averaged across five sessions. The response rates and their distribution were similar in both D2R-overexpressing mice and control EGFP mice, with the highest mean response rates near the target of 24 s. For a quantitative analysis of peak trial performance, we fit a Gaussian probability density function to peak trial data from individual mice, as previously done [45,53].

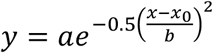

We found no significant differences in best-fit parameter values for peak location (*x*_*0*_) or peak width (*b*) suggesting no D2R-mediated alterations in the accuracy and precision in timing 24-s intervals **(Figures 4E**,**F)**. Moreover, we did not find alterations in maximal response rate estimates reflected in the peak height parameter (*a*) (**Figure 4G**), suggesting that motivation was not affected, consistent with our previous findings in a progressive ratio task [42]. These results, together with the temporal discrimination data, suggest that timing is not fundamentally altered by increased D2R expression levels in NAc CINs.

## DISCUSSION

Using Cre-mediated recombination with AAV gene transfer, we have found that selectively overexpressing D2Rs in CINs of the adult NAc leads to a significant increase in impulsive choice as measured in a delay discounting task. In line with these results, CIN-D2KO mice, which lack the *Drd2* gene in CINs, showed *decreased* delay discounting. These effects were not associated with alterations in sensitivity to reward magnitude or in the ability to time intervals.

Moreover, the behavioral impact of CIN D2Rs was not observed in probabilistic discounting, a related form of impulsive choice. Together, these findings indicate that D2R expression levels in CINs powerfully regulate impulsive decision-making processes involving delay costs.

Brain imaging and autoradiography studies have reported a correlation between lower D2R availability in ventral striatum, especially the NAc core, and higher trait motor and choice impulsivity in rats [21,22]. Whether alterations in NAc D2R levels or function cause impulsive choice behavior, however, has been difficult to demonstrate conclusively. Many of the studies to date have involved acute systemic administration of D2R/D3R pharmacological agents. Some report increased sensitivity to delay costs with D2R antagonism [20,54], whereas others show no effect with either antagonists or agonists [2,55,56]. Such discrepancies may relate, to some extent, to the combined impact of these agents on NAc and relevant extrastriatal regions (e.g., prefrontal cortex, amygdala, and ventral tegmental area), whose D2R signaling may have varied roles in impulsive decision-making [57-61]. However, the few studies that have performed intra-NAc microinfusions of raclopride, eticlopride or quinpirole have shown no effect of NAc D2R in delay discounting[62,63]. It is conceivable that even within the NAc, concurrent blockade of D2Rs in different cell types may mask their unique contributions. For example, shRNA-mediated knockdown of D2Rs in the VTA increases delay discounting[58]. While VTA dopamine neurons have brain-wide projections, it is plausible that presynaptic D2Rs in dopaminergic afferents to the NAc play a role in limiting impulsive decision-making. In contrast, our findings clearly demonstrate that D2Rs in NAc CINs increase delay discounting. This new information not only suggests that D2Rs expressed by different neuronal populations operating within the NAc can have opposing effects on impulsive choice, but also highlights the importance of cell-targeted strategies in unraveling dopamine’s complex modulation of NAc circuitry.

Growing evidence has also implicated cholinergic neurotransmission in delay discounting. Chronic smokers show greater impulsive choice, suggesting a permissive role for nicotinic acetylcholine receptor (nAChR) function [64,65]. Likewise, acute nicotine administration in rats leads to enhanced choice impulsivity, an effect that is prevented by nAChR blockade [36,37]. However, conflicting results have also been described [38,39]. Because these studies also relied on systemic delivery, the specific brain areas and cell types that are critically involved remain to be defined. The behavioral effects that we observed following selective targeting of NAc CINs clearly identify these neurons as a key node in the neurocircuitry underlying impulsive choice.

Performance in both probabilistic discounting and delay discounting tasks depends on an intact NAc core and can be sensitive to D2R blockade or nicotine [11,20,39,66,67]. However, the manipulations of CIN D2Rs in our study did not affect probabilistic discounting, indicating that these receptors are not broadly involved in all forms of impulsive choice, but selectively contribute to decision-making involving delay costs. Although intra-NAc delivery of D2R agonist increases risk-seeking behavior, biasing choice toward larger, probabilistic rewards[68], it is possible that D2R-expressing cell types in the NAc other than CINs play a more prominent role in this behavior. Supporting this hypothesis is the fact that phasic activity in D2R-expressing spiny projection neurons (SPNs) is sufficient to decrease risk preference[68]. Whether D2Rs in SPNs mediate probabilistic discounting would need to be directly tested.

Altered sensitivity to reward magnitude can result in impulsive choice if an animal is unable perceive a large reward as more valuable than a small reward. Neither of our CIN D2Rs manipulations affected discrimination between the large and small reinforcers in sessions involving no delay or probability costs. These findings suggest that CIN D2Rs do not contribute to the independent processing of reward magnitude, in agreement with several studies using NAc lesions and systemic D2R or cholinergic receptor drugs [11,36,38,63,67,69,70].

The ability to accurately represent the time interval between a reward-seeking action and reward retrieval is intricately linked to delay intolerance. For example, overestimation of elapsed time could reduce preference for the larger delayed reward[71]. Indeed, individuals deemed impulsive on delay discounting tasks are more prone to timing errors compared to control subjects [72]. Similarly, rats that showed higher timing precision in peak interval and temporal discrimination tasks also show reduced delay-based impulsivity [52,73]. Multiple studies also support a role of dopamine in timing [74-76]. Transgenic mice that selectively overexpress D2Rs in striatal SPNs since early in development show reduced timing precision in a peak interval task as well as deficits in timing long sample durations in a temporal discrimination paradigm[44,45]. In contrast, we did not observe an effect of CIN D2R upregulation in either timing task, suggesting that the effect of CIN D2Rs on delay discounting does not involve alterations in representation of time.

Choices made in delay discounting tasks require a dynamic, subjective assessment of reward value that integrates the magnitude and the changing delay properties of a reward [9]. The NAc appears to be a key site for this integration across species. In humans, neural activity in ventral striatum during delay discounting is more strongly correlated with subjective value than to objective reward characteristics like magnitude and delay [9]. Furthermore, inactivation of the NAc core in rats decreased discounting only in a task that measured sensitivity to both delay and magnitude but has no effect when these factors were independently adjusted [13].

The cellular mechanisms underlying integration of these reward characteristics in intertemporal choice, however, are not well understood, but it may involve distinct subsets of striatal neurons whose activity is modulated by both reward size and delay[77-79]. Further, cue-evoked activity of a subset of neurons in the dorsal caudate nucleus encodes the temporally discounted value of rewards but not reward magnitude or delay alone [78]. Whether a similar dynamic computation of subjective value occurs in specific subset(s) of NAc neurons during delay discounting tasks remains to be determined, but recent evidence indicates that NAc dopamine may contribute to this process. For instance, cue-evoked dopamine release in NAc not only encodes the relative value of small and large reward options, but also how that value changes with increasing delays[19]. Because cue-evoked dopamine and acetylcholine signals temporally coincide in mouse striatum [34,35], it is tempting to speculate that D2R-dependent modulation of NAc CINs contributes to integration of reward size and delay information in impulsive decision-making.

Growing evidence suggests that discounting of delayed rewards is a stable, heritable trait that contributes to the etiology and treatment outcomes of various mental health disorders[3,8,80]. Despite the important clinical implications, few pharmacological interventions are currently available that are specific to impulsivity and that are based on an understanding of its complex subdomains and underlying neurocircuitry. Our findings identify NAc CIN D2Rs as critical players in the mechanisms of delay-based impulsive choice. This new information refines our current understanding of the contributions of striatal dopamine and acetylcholine to impulsive behavior and raise the possibility that modulation of NAc acetylcholine might hold promise for more targeted treatments for choice impulsivity.

## ACKNOWLEDGMENTS

We thank David D’Onofrio, Eric Teboul, Emily Huegler, and Daphne Baker for technical assistance, Katherine Nautiyal for input on behavioral design and analysis, and Christoph Kellendonk and Jonathan Javitch for sharing the D2R viral vector.

## AUTHOR CONTRIBUTIONS

J.C., J.Y., J.F. and B.A. and E.F.G. conducted the experiments and data analysis. E.F.G and J.C. wrote the manuscript. P.D. and B.A. edited the manuscript. E.F.G. and P.D. designed the experiments. E.F.G. supervised the experiments and data analysis.

## REFERENCES

1 Ainslie G. Specious reward: A behavioral theory of impulsiveness and impulse control. Psychol Bull. 1975;82:463–96.

2 Evenden JL, Ryan CN. The pharmacology of impulsive behaviour in rats: the effects of drugs on response choice with varying delays of reinforcement. Psychopharmacology (Berl). 1996;128(2):161–70.

3 Bickel WK, Jarmolowicz DP, Mueller ET, Koffarnus MN, Gatchalian KM. Excessive discounting of delayed reinforcers as a trans-disease process contributing to addiction and other disease-related vulnerabilities: Emerging evidence. Pharmacol Ther. 2012;134(3):287–97.

4 Patros CH, Alderson RM, Kasper LJ, Tarle SJ, Lea SE, Hudec KL. Choice-impulsivity in children and adolescents with attention-deficit/hyperactivity disorder (ADHD): A meta-analytic review. Clin Psychol Rev. 2016;43:162–74.

5 Heerey EA, Robinson BM, McMahon RP, Gold JM. Delay discounting in schizophrenia. Cogn Neuropsychiatry. 2007;12(3):213–21.

6 Weller RE, Cook EW, Avsar KB, Cox JE. Obese women show greater delay discounting than healthy-weight women. Appetite. 2008;51(3):563–69.

7 Daugherty JR, Brase GL. Taking time to be healthy: Predicting health behaviors with delay discounting and time perspective. Pers Individ Dif. 2010;48(2):202–07.

8 Lempert KM, Steinglass JE, Pinto A, Kable JW, Simpson HB. Can delay discounting deliver on the promise of RDoC? Psychol Med. 2019;49(2):190–99.

9 Kable JW, Glimcher PW. The neural correlates of subjective value during intertemporal choice. Nat Neurosci. 2007;10(12):1625–33.

10 Lempert KM, Speer ME, Delgado MR, Phelps EA. Positive autobiographical memory retrieval reduces temporal discounting. Soc Cogn Affect Neurosci. 2017;12(10):1584–93.

11 Cardinal RN, Pennicott DR, Sugathapala CL, Robbins TW, Everitt BJ. Impulsive choice induced in rats by lesions of the nucleus accumbens core. Science. 2001;292(5526):2499–501.

12 da Costa Araújo S, Body S, Hampson CL, Langley RW, Deakin JF, Anderson IM, et al. Effects of lesions of the nucleus accumbens core on inter-temporal choice: further observations with an adjusting-delay procedure. Behav Brain Res. 2009;202(2):272–7.

13 Moschak TM, Mitchell SH. Partial inactivation of nucleus accumbens core decreases delay discounting in rats without affecting sensitivity to delay or magnitude. Behav Brain Res. 2014;268:159–68.

14 Roesch MR, Calu DJ, Schoenbaum G. Dopamine neurons encode the better option in rats deciding between differently delayed or sized rewards. Nat Neurosci. 2007;10(12):1615–24.

15 Kobayashi S, Schultz W. Influence of Reward Delays on Responses of Dopamine Neurons. The Journal of Neuroscience. 2008;28(31):7837–46.

16 Joutsa J, Voon V, Johansson J, Niemelä S, Bergman J, Kaasinen V. Dopaminergic function and intertemporal choice. Translational Psychiatry. 2015;5(1):e491–e91.

17 Winstanley CA, Theobald DE, Dalley JW, Robbins TW. Interactions between serotonin and dopamine in the control of impulsive choice in rats: therapeutic implications for impulse control disorders. Neuropsychopharmacology. 2005;30(4):669–82.

18 Day JJ, Jones JL, Wightman RM, Carelli RM. Phasic Nucleus Accumbens Dopamine Release Encodes Effort- and Delay-Related Costs. Biol Psychiatry. 2010;68(3):306–09.

19 Saddoris MP, Sugam JA, Stuber GD, Witten IB, Deisseroth K, Carelli RM. Mesolimbic Dopamine Dynamically Tracks, and Is Causally Linked to, Discrete Aspects of Value-Based Decision Making. Biol Psychiatry. 2015;77(10):903–11.

20 Wade TR, de Wit H, Richards JB. Effects of dopaminergic drugs on delayed reward as a measure of impulsive behavior in rats. Psychopharmacology (Berl). 2000;150(1):90–101.

21 Dalley JW, Fryer TD, Brichard L, Robinson ESJ, Theobald DEH, Lääne K, et al. Nucleus Accumbens D2/3 Receptors Predict Trait Impulsivity and Cocaine Reinforcement. Science. 2007;315(5816):1267–70.

22 Barlow RL, Gorges M, Wearn A, Niessen HG, Kassubek J, Dalley JW, et al. Ventral Striatal D2/3 Receptor Availability Is Associated with Impulsive Choice Behavior As Well As Limbic Corticostriatal Connectivity. Int J Neuropsychopharmacol. 2018;21(7):705–15.

23 Gallo EF. Disentangling the diverse roles of dopamine D2 receptors in striatal function and behavior. Neurochem Int. 2019;125:35–46.

24 Bolam JP, Wainer BH, Smith AD. Characterization of cholinergic neurons in the rat neostriatum. A combination of choline acetyltransferase immunocytochemistry, Golgi-impregnation and electron microscopy. Neuroscience. 1984;12(3):711–8.

25 Matamales M, Gotz J, Bertran-Gonzalez J. Quantitative Imaging of Cholinergic Interneurons Reveals a Distinctive Spatial Organization and a Functional Gradient across the Mouse Striatum. PLoS One. 2016;11(6):e0157682.

26 Calabresi P, Centonze D, Gubellini P, Pisani A, Bernardi G. Blockade of M2-like muscarinic receptors enhances long-term potentiation at corticostriatal synapses. Eur J Neurosci. 1998;10(9):3020–3.

27 Wang Z, Kai L, Day M, Ronesi J, Yin HH, Ding J, et al. Dopaminergic control of corticostriatal long-term synaptic depression in medium spiny neurons is mediated by cholinergic interneurons. Neuron. 2006;50(3):443–52.

28 Cui G, Jun SB, Jin X, Pham MD, Vogel SS, Lovinger DM, et al. Concurrent activation of striatal direct and indirect pathways during action initiation. Nature. 2013;494(7436):238–42.

29 Soares-Cunha C, de Vasconcelos NAP, Coimbra B, Domingues AV, Silva JM, Loureiro-Campos E, et al. Nucleus accumbens medium spiny neurons subtypes signal both reward and aversion. Mol Psychiatry. 2020;25(12):3241–55.

30 Threlfell S, Lalic T, Platt NJ, Jennings KA, Deisseroth K, Cragg SJ. Striatal dopamine release is triggered by synchronized activity in cholinergic interneurons. Neuron. 2012;75(1):58–64.

31 Cachope R, Mateo Y, Mathur BN, Irving J, Wang HL, Morales M, et al. Selective activation of cholinergic interneurons enhances accumbal phasic dopamine release: setting the tone for reward processing. Cell reports. 2012;2(1):33–41.

32 Liu C, Cai X, Ritzau-Jost A, Kramer PF, Li Y, Khaliq ZM, et al. An action potential initiation mechanism in distal axons for the control of dopamine release. Science. 2022;375(6587):1378–85.

33 Aosaki T, Graybiel AM, Kimura M. Effect of the nigrostriatal dopamine system on acquired neural responses in the striatum of behaving monkeys. Science. 1994;265(5170):412–5.

34 Gallo EF, Greenwald J, Yeisley J, Teboul E, Martyniuk KM, Villarin JM, et al. Dopamine D2 receptors modulate the cholinergic pause and inhibitory learning. Mol Psychiatry. 2022;27(3):1502–14.

35 Martyniuk KM, Torres-Herraez A, Lowes DC, Rubinstein M, Labouesse MA, Kellendonk C. Dopamine D2Rs coordinate cue-evoked changes in striatal acetylcholine levels. eLife. 2022;11:e76111.

36 Kolokotroni KZ, Rodgers RJ, Harrison AA. Acute nicotine increases both impulsive choice and behavioural disinhibition in rats. Psychopharmacology (Berl). 2011;217(4):455–73.

37 Dallery J, Locey ML. Effects of acute and chronic nicotine on impulsive choice in rats. Behav Pharmacol. 2005;16(1):15–23.

38 Ozga JE, Anderson KG. Reduction in delay discounting due to nicotine and its attenuation by cholinergic antagonists in Lewis and Fischer 344 rats. Psychopharmacology (Berl). 2018;235(1):155–68.

39 Mendez IA, Gilbert RJ, Bizon JL, Setlow B. Effects of acute administration of nicotinic and muscarinic cholinergic agonists and antagonists on performance in different cost-benefit decision making tasks in rats. Psychopharmacology (Berl). 2012;224(4):489–99.

40 Gong S, Doughty M, Harbaugh CR, Cummins A, Hatten ME, Heintz N, et al. Targeting Cre Recombinase to Specific Neuron Populations with Bacterial Artificial Chromosome Constructs. The Journal of Neuroscience. 2007;27(37):9817–23.

41 Bello EP, Mateo Y, Gelman DM, Noain D, Shin JH, Low MJ, et al. Cocaine supersensitivity and enhanced motivation for reward in mice lacking dopamine D2 autoreceptors. Nat Neurosci. 2011;14(8):1033–8.

42 Gallo EF, Meszaros J, Sherman JD, Chohan MO, Teboul E, Choi CS, et al. Accumbens dopamine D2 receptors increase motivation by decreasing inhibitory transmission to the ventral pallidum. Nat Commun. 2018;9(1):1086.

43 Nautiyal KM, Wall MM, Wang S, Magalong VM, Ahmari SE, Balsam PD, et al. Genetic and Modeling Approaches Reveal Distinct Components of Impulsive Behavior. Neuropsychopharmacology. 2017;42(6):1182–91.

44 Ward RD, Kellendonk C, Simpson EH, Lipatova O, Drew MR, Fairhurst S, et al. Impaired timing precision produced by striatal D2 receptor overexpression is mediated by cognitive and motivational deficits. Behav Neurosci. 2009;123(4):720–30.

45 Drew MR, Simpson EH, Kellendonk C, Herzberg WG, Lipatova O, Fairhurst S, et al. Transient overexpression of striatal D2 receptors impairs operant motivation and interval timing. J Neurosci. 2007;27(29):7731–9.

46 Pinkston JW, Lamb RJ. Delay discounting in C57BL/6J and DBA/2J mice: adolescent-limited and life-persistent patterns of impulsivity. Behav Neurosci. 2011;125(2):194–201.

47 Cardinal RN. Neural systems implicated in delayed and probabilistic reinforcement. Neural Netw. 2006;19(8):1277–301.

48 Richards JB, Zhang L, Mitchell SH, de Wit H. Delay or probability discounting in a model of impulsive behavior: effect of alcohol. J Exp Anal Behav. 1999;71(2):121–43.

49 Kharkwal G, Brami-Cherrier K, Lizardi-Ortiz JE, Nelson AB, Ramos M, Del Barrio D, et al. Parkinsonism Driven by Antipsychotics Originates from Dopaminergic Control of Striatal Cholinergic Interneurons. Neuron. 2016;91(1):67–78.

50 Augustin SM, Chancey JH, Lovinger DM. Dual Dopaminergic Regulation of Corticostriatal Plasticity by Cholinergic Interneurons and Indirect Pathway Medium Spiny Neurons. Cell reports. 2018;24(11):2883–93.

51 Lee JH, Ribeiro EA, Kim J, Ko B, Kronman H, Jeong YH, et al. Dopaminergic Regulation of Nucleus Accumbens Cholinergic Interneurons Demarcates Susceptibility to Cocaine Addiction. Biol Psychiatry. 2020;88(10):746–57.

52 McClure J, Podos J, Richardson H. Isolating the delay component of impulsive choice in adolescent rats. Front Integr Neurosci. 2014;8(3).

53 Buhusi CV, Meck WH. Timing for the absence of a stimulus: the gap paradigm reversed. J Exp Psychol Anim Behav Process. 2000;26(3):305–22.

54 Denk F, Walton ME, Jennings KA, Sharp T, Rushworth MF, Bannerman DM. Differential involvement of serotonin and dopamine systems in cost-benefit decisions about delay or effort. Psychopharmacology (Berl). 2005;179(3):587–96.

55 Koffarnus MN, Newman AH, Grundt P, Rice KC, Woods JH. Effects of selective dopaminergic compounds on a delay-discounting task. Behav Pharmacol. 2011;22(4):300–11.

56 van Gaalen MM, van Koten R, Schoffelmeer AN, Vanderschuren LJ. Critical involvement of dopaminergic neurotransmission in impulsive decision making. Biol Psychiatry. 2006;60(1):66–73.

57 Winstanley CA, Theobald DE, Cardinal RN, Robbins TW. Contrasting roles of basolateral amygdala and orbitofrontal cortex in impulsive choice. J Neurosci. 2004;24(20):4718–22.

58 Bernosky-Smith KA, Qiu YY, Feja M, Lee YB, Loughlin B, Li JX, et al. Ventral tegmental area D2 receptor knockdown enhances choice impulsivity in a delay-discounting task in rats. Behav Brain Res. 2018;341:129–34.

59 Zeeb FD, Floresco SB, Winstanley CA. Contributions of the orbitofrontal cortex to impulsive choice: interactions with basal levels of impulsivity, dopamine signalling, and reward-related cues. Psychopharmacology (Berl). 2010;211(1):87–98.

60 Pardey MC, Kumar NN, Goodchild AK, Cornish JL. Catecholamine receptors differentially mediate impulsive choice in the medial prefrontal and orbitofrontal cortex. Journal of Psychopharmacology. 2013;27(2):203–12.

61 Larkin JD, Jenni NL, Floresco SB. Modulation of risk/reward decision making by dopaminergic transmission within the basolateral amygdala. Psychopharmacology (Berl). 2016;233(1):121–36.

62 Yates JR, Bardo MT. Effects of intra-accumbal administration of dopamine and ionotropic glutamate receptor drugs on delay discounting performance in rats. Behav Neurosci. 2017;131(5):392–405.

63 Li Y, Zuo Y, Yu P, Ping X, Cui C. Role of basolateral amygdala dopamine D2 receptors in impulsive choice in acute cocaine-treated rats. Behav Brain Res. 2015;287:187–95.

64 Field M, Santarcangelo M, Sumnall H, Goudie A, Cole J. Delay discounting and the behavioural economics of cigarette purchases in smokers: the effects of nicotine deprivation. Psychopharmacology (Berl). 2006;186(2):255–63.

65 Bickel WK, Odum AL, Madden GJ. Impulsivity and cigarette smoking: delay discounting in current, never, and ex-smokers. Psychopharmacology (Berl). 1999;146(4):447–54.

66 St Onge JR, Floresco SB. Dopaminergic modulation of risk-based decision making. Neuropsychopharmacology. 2009;34(3):681–97.

67 Cardinal RN, Howes NJ. Effects of lesions of the nucleus accumbens core on choice between small certain rewards and large uncertain rewards in rats. BMC Neurosci. 2005;6:37.

68 Zalocusky KA, Ramakrishnan C, Lerner TN, Davidson TJ, Knutson B, Deisseroth K. Nucleus accumbens D2R cells signal prior outcomes and control risky decision-making. Nature. 2016;531(7596):642–6.

69 Steele CC, Peterson JR, Marshall AT, Stuebing SL, Kirkpatrick K. Nucleus accumbens core lesions induce sub-optimal choice and reduce sensitivity to magnitude and delay in impulsive choice tasks. Behav Brain Res. 2018;339:28–38.

70 Balleine B, Killcross S. Effects of ibotenic acid lesions of the Nucleus Accumbens on instrumental action. Behav Brain Res. 1994;65(2):181–93.

71 Wittmann M, Paulus MP. Decision making, impulsivity and time perception. Trends Cogn Sci. 2008;12(1):7–12.

72 Baumann AA, Odum AL. Impulsivity, risk taking, and timing. Behav Processes. 2012;90(3):408–14.

73 Marshall AT, Smith AP, Kirkpatrick K. Mechanisms of impulsive choice: I. Individual differences in interval timing and reward processing. J Exp Anal Behav. 2014;102(1):86–101.

74 Meck WH. Neuroanatomical localization of an internal clock: A functional link between mesolimbic, nigrostriatal, and mesocortical dopaminergic systems. Brain Res. 2006;1109(1):93–107.

75 Buhusi CV, Meck WH. Differential effects of methamphetamine and haloperidol on the control of an internal clock. Behav Neurosci. 2002;116(2):291–7.

76 Balci F, Ludvig EA, Abner R, Zhuang X, Poon P, Brunner D. Motivational effects on interval timing in dopamine transporter (DAT) knockdown mice. Brain Res. 2010;1325:89–99.

77 Roesch M, Bryden D. Impact of Size and Delay on Neural Activity in the Rat Limbic Corticostriatal System. Front Neurosci. 2011;5.

78 Hori Y, Mimura K, Nagai Y, Fujimoto A, Oyama K, Kikuchi E, et al. Single caudate neurons encode temporally discounted value for formulating motivation for action. eLife. 2021;10:e61248.

79 Cai X, Kim S, Lee D. Heterogeneous Coding of Temporally Discounted Values in the Dorsal and Ventral Striatum during Intertemporal Choice. Neuron. 2011;69(1):170–82.

80 Anokhin AP, Grant JD, Mulligan RC, Heath AC. The genetics of impulsivity: evidence for the heritability of delay discounting. Biol Psychiatry. 2015;77(10):887–94.

